# Dynamic visualisation of million-tip trees: the OneZoom project

**DOI:** 10.1101/2020.10.14.323055

**Authors:** Yan Wong, James Rosindell

**Affiliations:** OneZoom CIO, Office 7, 35-37 Ludgate Hill, London, EC4M 7JN, UK; Big Data Institute, University of Oxford, Oxford, UK; Department of Life Sciences, Silwood Park Campus, Imperial College London, Buckhurst Road, Ascot, Berkshire, SL5 7PY, UK

## Abstract

1. The complete tree of life is now available, but methods to visualise it are still needed to meet needs in research, teaching and science communication. Dynamic visualisation of million-tip trees requires many challenges in data synthesis, data handling and computer graphics to be overcome.
2. Our approach is to automate data processing, synthesise data from a wide range of available sources, then to feed these data to a client-side visualisation engine in parts. We develop a way to store the whole tree topology locally in a highly compressed form, then dynamically populate metadata such as text and images as the user explores.
3. The result is a seamless and smooth way to explore the complete tree of life, including images and metadata, even on relatively old mobile devices.
4. The underlying methods developed have applications that transcend tree of life visualisation. For the whole complete tree, we describe automated ID mappings between well known resources without resorting to taxonomic name resolution, automated methods to collate sets of public domain representative images for higher taxa, and an index to measure public interest of individual species.
5. The visualisation layout and the client user interface are both abstracted components of the codebase enabling other zoomable tree layouts to be swapped in, and supporting multiple applications including exhibition kiosks and digital art.
6. After ten years of work, our tree of life explorer is now broadly complete, it has attracted nearly 1.5 million online users, and is backed by a novel long-term sustainability plan. We conclude our description of the OneZoom project by suggesting the next challenges that need to be solved in this field: extinct species and guided tours around the tree.

## Introduction

Over the past decade phylogenetic datasets have expanded in size, and evolutionary trees with many millions of tips are now available (Hinchliff et al. 2015). Methods to visualise the complete tree of life are needed for research, teaching, and science communication. However, standard tree drawing tools are inappropriate for visualizing such huge trees (Page 2012; James Rosindell and Wong 2018): static images do not provide enough resolution, and the need for extreme precision when zooming into highly nested trees makes many off-the-shelf interactive tools unsuitable.

We previously described proof of concept visualization software for a large tree explorer with a deep zooming user interface, enabling web browsers to render arbitrarily large phylogenetic trees by using a nested, fractal layout while dynamically re-anchoring the coordinate system to remove precision errors (J. Rosindell and Harmon 2012). However, the complete tree of life, comprising many millions of species together with names and images, constitutes gigabytes of information: too much to provide in a single statically-loaded page.

Here, we describe a new client-server software package, including numerous improvements, that mostly completes the plan of work laid out in our original concept. This new software allows a phylogeny of all species of life on earth to be visualised on a single web page, with fast initial loading as well as completely smooth navigation around the tree, even over a relatively slow internet connection and on mobile devices. It also supports an expandable range of different visualisation layouts.

Our software provides automated methods to embellish the tree with essential metadata: not only scientific and vernacular names (in multiple languages), but also images, extinction risk (IUCN 2020), and links to online resources such as Wikipedia, the Encyclopedia of Life (Parr et al. 2014), NCBI/GenBank (NCBI Resource Coordinators 2018) and GBIF (GBIF 2021). To link metadata to millions of tree nodes, we develop code that maps nodes to permanent identifiers in other biodiversity databases; additional scripts use these mappings to download appropriate metadata from external sources. As well as enabling annotation of the tree, such mappings also enable synthesis and analysis (Thessen et al. 2018). For example, we can automatically produce a set of high quality, phylogenetically diverse images to represent higher taxonomic groups, and construct a “popularity index”, measuring the public prominence of every species on the tree.

Typically, a biodiversity browser is a database of taxa pages with some taxonomic organisation given primarily as a way to navigate between the pages. Our project is different in two respects, first, the organisation is itself the main focus and second our organisation is based on phylogeny, not taxonomy. Taxonomic classifications, such as those provided by NCBI, may include non-monophyletic groups such as “fish” or “bacteria”, and indeed many genera are also non-monophyletic. In contrast, non-monophyletic names of any kind have no natural place on a phylogenetic tree. Another distinction is that even when strictly hierarchical, taxonomies are usually coarsely resolved such that a single node may have a large number of immediate children; while taxonomists do strive to find order in nodes with huge numbers of children, and do so in a phylogenetic manner, the optimal end point is a functional categorisation which might be to have 20-30 children in each category. In contrast, phylogenetic trees reflect evolutionary history alone: truly multifurcating parent nodes (polytomies) are much less common, and almost always reflect uncertainty. So, the optimal end point is a bifurcating tree reflecting evolutionary history. Focussing on phylogeny restricts the potential data sources for the tree: indeed we are aware of only one attempt at synthesising a phylogeny of all life, the Open Tree of Life (Hinchliff et al. 2015), henceforth referred to as the OpenTree. This open dataset is therefore the primary data source for the OneZoom Tree of Life Explorer (hereafter just our “ToL explorer”) that we make freely available at http://www.onezoom.org (OneZoom Core Team 2021a).

The source code to our visualizer and associated server software is freely downloadable from https://github.com/OneZoom/ and the version 3.5 ‘Chocolate chip starfish’ to accompany this manuscript is also available in a Docker container (OneZoom Core Team 2021b). To the best of our knowledge, our ToL explorer is the only seamless visualization software that displays the complete tree of described life, the only complete tree of life explorer incorporating images and other metadata, and provides the only popularity index covering all described species on this planet.

## Materials and Methods

Our ToL explorer is best understood in three layers. *External data* (1) is collected from multiple sources, and undergoes *server side processing* (2), resulting in an online API which is accessed by a *client side tree viewer* (3). The tree viewer is written in ECMAscript, compiled to Javascript, and thus usable in any modern web browser; the API is served using the Web2Py framework (Di Pierro 2013) and hence is primarily written in Python v3. The API can be also used by other unrelated projects to access derived data products, such as taxon popularity. Figure 1 provides a schematic, the components of which are described in more detail below.

**Figure 1:**
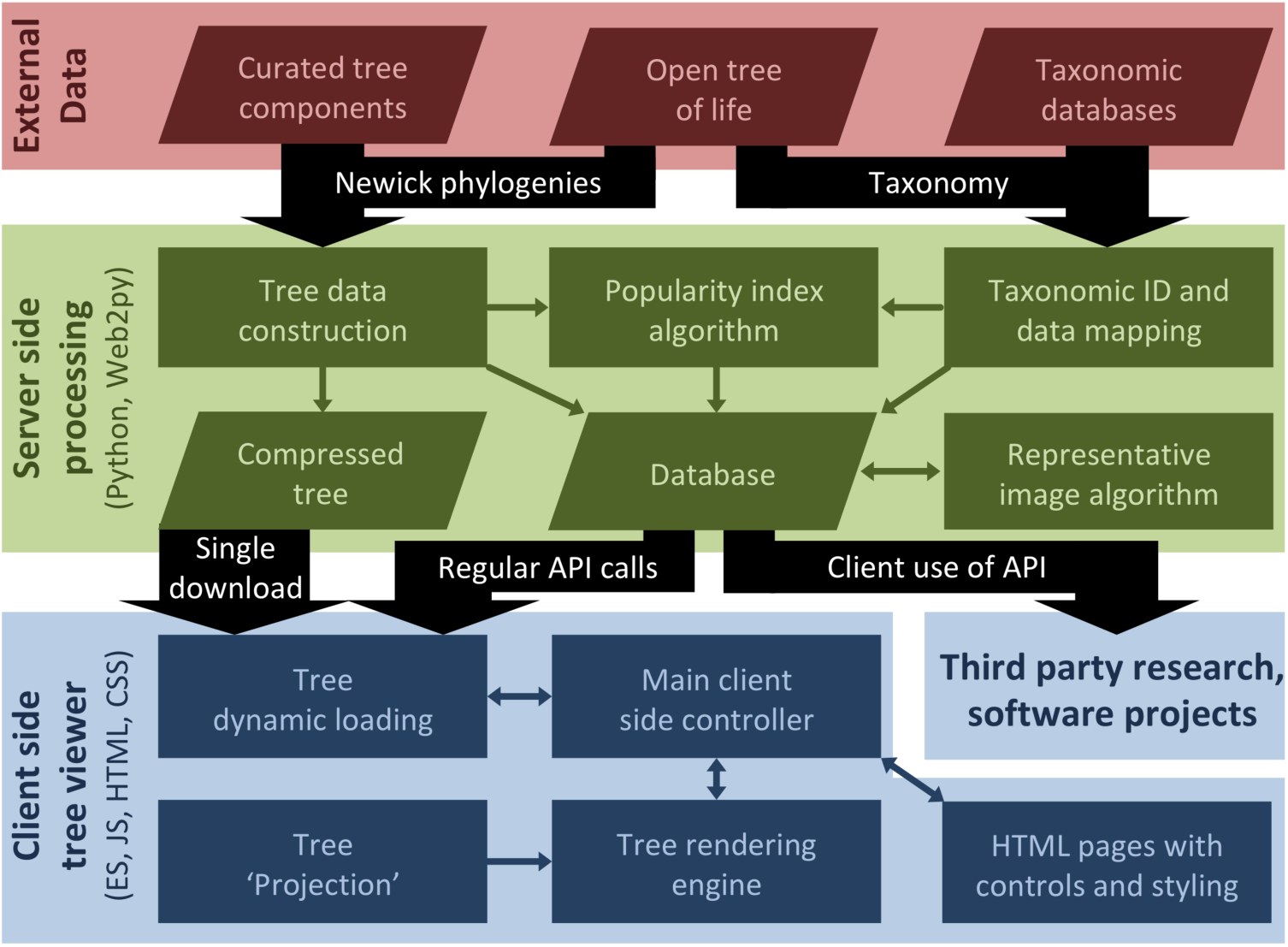
Overview schematic of all of the key ToL explorer components. The main three layers of the system, external data, server side processing, and client side processing are shown in different colours, with annotated connections between them. Components within a layer consist of data products (slanted parallelograms) or algorithms and other code (rectangles), with connections between them indicated using small arrows. The six server side processing blocks, together with the client side tree viewer layer, are described in seven respective subsections of the methods.

### Tree data construction

We use the most recent available “synthetic tree” from the OpenTree project as our core data source, trimming the data before final output to remove divisions below the species level. However, the synthetic OpenTree does not currently contain node date estimates, and we deem it inaccurate in various places, such as in its support for a monophyletic Archaea. We have therefore developed a set of Python scripts and configuration files which automatically extract large subtrees from the OpenTree topology and graft them onto a hand-curated backbone (see supporting information S3). This onerous step should become largely redundant as the OpenTree improves in backbone accuracy and starts incorporating dates (Hinchliff et al. 2015).

### Compressed tree

At the heart of the OneZoom approach is the realisation that, unlike geographical maps, hierarchical relationships can be represented by regular geometric structures that require very little information to encode. In particular, topological relationships between taxa have a compact encoding that, when coupled with a visualisation method, uniquely determines the geometry of a phylogenetic tree. Hence the entire topology can be downloaded before visualization, and rendered into 2D space in the client’s browser. Metadata such as names and images can then be asynchronously layered over the visualisation using an online API. This results in smooth zooming and navigation, in contrast to the substantially more bandwidth-intensive approach of downloading pre-computed tiled images, as used by most online geographical mapping software and extended to biological trees by de Vienne (2016).

To compress the tree topology we represent it as an unlabelled string of nested parentheses similar to the well-known Newick format. We create a single bifurcating tree by randomly resolving all polytomies, then “ladderizing” it, with the result that the commas conventionally used to separate taxa are redundant, and can be omitted (see Figure 2). The arbitrary splits created by polytomy resolution are marked using curly versus round parentheses; this allows the client to determine which internal nodes are “real” and which are “fake” nodes introduced by polytomy splitting. The client can index leaves and internal nodes by incremental counting along the string: these temporary indexes map to the row number in two associated SQL database tables. The server scripts used to regularly update the phylogeny must therefore jointly create a new compressed tree string and rebuild these two tables (supporting information S4).

**Figure 2:**
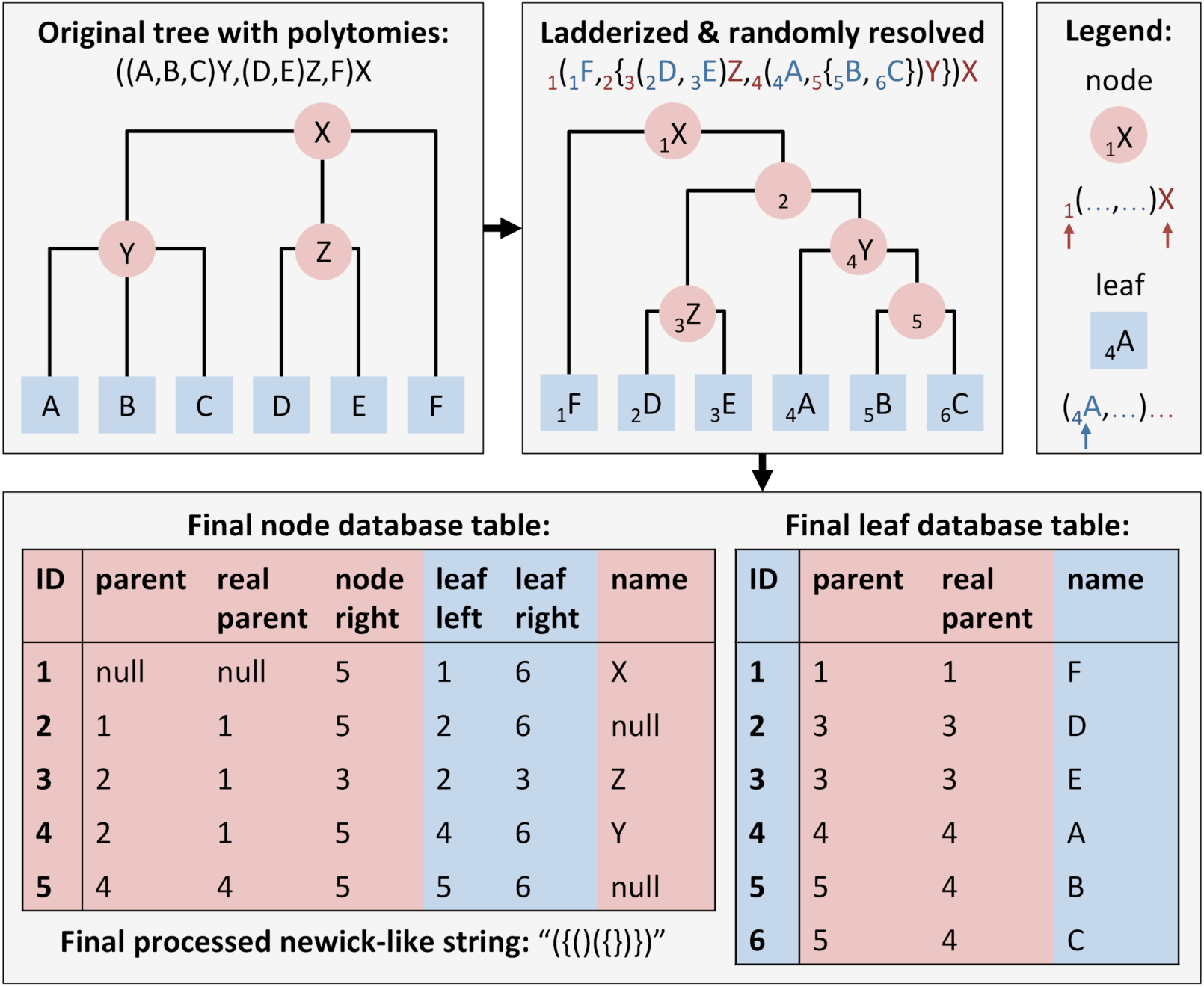
Example tree showing initial tree in classic newick format, followed by the same tree, now with randomly resolved polytomies. Finally we show the resulting database tables and processed newick string ready for fast navigation. Note that in this figure, we omit extra fields in the leaf and node tables, such as OTT ID, as they are unrelated to nesting and indexing.

The current 2.2 million species on our tree produce a text string of 4.4 million characters in length, compressed by gzip to 660kb (roughly 2.5 bits per species). While this is quick to download, client-side handling of such a long string can lead to performance issues. To address this, we also provide a precalculated index of “cutpoints” giving the start and end indexes of substrings representing large monophyletic clades. This way the client never has to scan through the entire string to match distant opening and closing brackets (see Figure 2).

### Taxonomic ID and data mapping

Leaf and node indexes in the compressed tree string will change with each updating of the phylogeny, so each taxon on the tree also needs a unique identifier which is robust to changes in tree structure. Historically, the scientific name serves this function, but it becomes problematic for whole-life datasets (Page 2008): not only can plants and animals share identical names (e.g. *Ficus variegata*), but even within kingdoms, taxa at levels above the family are not required to be unique. Moreover names are subject to extensive synonymy and misspelling. As our project relies substantially on the OpenTree phylogeny, we use the “Open Tree Taxonomy” (OTT) ID as our canonical taxon identifier: this is a semi-permanent number assigned by the OpenTree project to virtually all leaves, and many internal nodes of the tree of life (Rees and Cranston 2017). The key principle we apply here is not to attempt taxonomic name resolution on the scientific name, as other resources already focus on such services (Boyle et al. 2013; Mozzherin, Myltsev, and Patterson 2017). Instead, the OTT ID gives alphanumerical identifiers into source datasets, such as those provided by NCBI (NCBI Resource Coordinators 2018) or GBIF (GBIF: The Global Biodiversity Information Facility 2020). We combine these to map to unique identifiers (UIDs) associated with additional online resources such as the Encyclopedia of Life (Parr et al. 2014), Wikidata (Vrandečić and Krötzsch 2014), and the IUCN Red List of Threatened Species (IUCN 2020), in a similar manner to Thessen *et al*. (2018) and allowing for inevitable inconsistencies due to likely errors in one or more source datasets (see supporting information S4).

### Representative image algorithm

Reliable mapping to Encyclopedia of Life (EOL) UIDs allows the OneZoom server to download images for over 80,000 leaves, and vernacular names in over 200 languages for approximately 191,000 taxa on the tree of life. Using the EOL public API (https://eol.org/docs/what-is-eol/classic-apis) OneZoom scripts saved up to 3 square thumbnail images per species, using user-provided crop points where historically provided by EOL. Each image has license, copyright holder, and source information, a set of quality ratings from EOL users, and whether the species has been verified as correctly identified (see supporting information S5). We download only images marked as Public Domain, Creative Commons AttributionOnly, or Creative Commons Attribution-ShareAlike, excluding images restricted to non-commercial use due lack of clarity on what is considered commercial. The 3 thumbnails include the best quality image together with an alternative public domain and verified image if required; this enables visualizations that consist of entirely public domain contributions. Initially, the scripts harvested over 100,000 images in a few weeks; they have previously been run continuously to update species images as more become available, however they are currently paused pending updates from EOL to reintroduce image quality ratings and optimal crop points. Cropped thumbnails are stored as UID-named jpg images on the OneZoom server filesystem, with the file name and relevant details stored in a database table; our system also allows for non-EOL-sourced images to be added by hand.

A novel postorder traversal algorithm was devised to produce a representative set of images summarising the biodiversity at internal nodes on the bifurcating tree (see supporting information S5). Eight images representing a parent node are selected so that if possible, and in order of priority: i) each child node contributes at least one image ii) all 8 slots for a parent are filled iii) the number of images contributed by each child aims to be proportional to the child’s species richness. The set of parent images starts with the first image from the richest child, then takes the first from the other child and so on in an alternating fashion until conditions (i)-(iii) mean that images are taken entirely from one child. Since the ordering of images determines which images get passed up the tree, once the set is filled we adjust its order by exchanging adjacent images if one has a substantially lower quality than the next in the set. This algorithm results in excellent quality, phylogenetically representative sets of images at all higher nodes in the tree (see Figure 3), and is run separately for best-quality, public domain, and verified images. The resulting representative image sets are available through an API (see supporting information S1).

**Figure 3:**
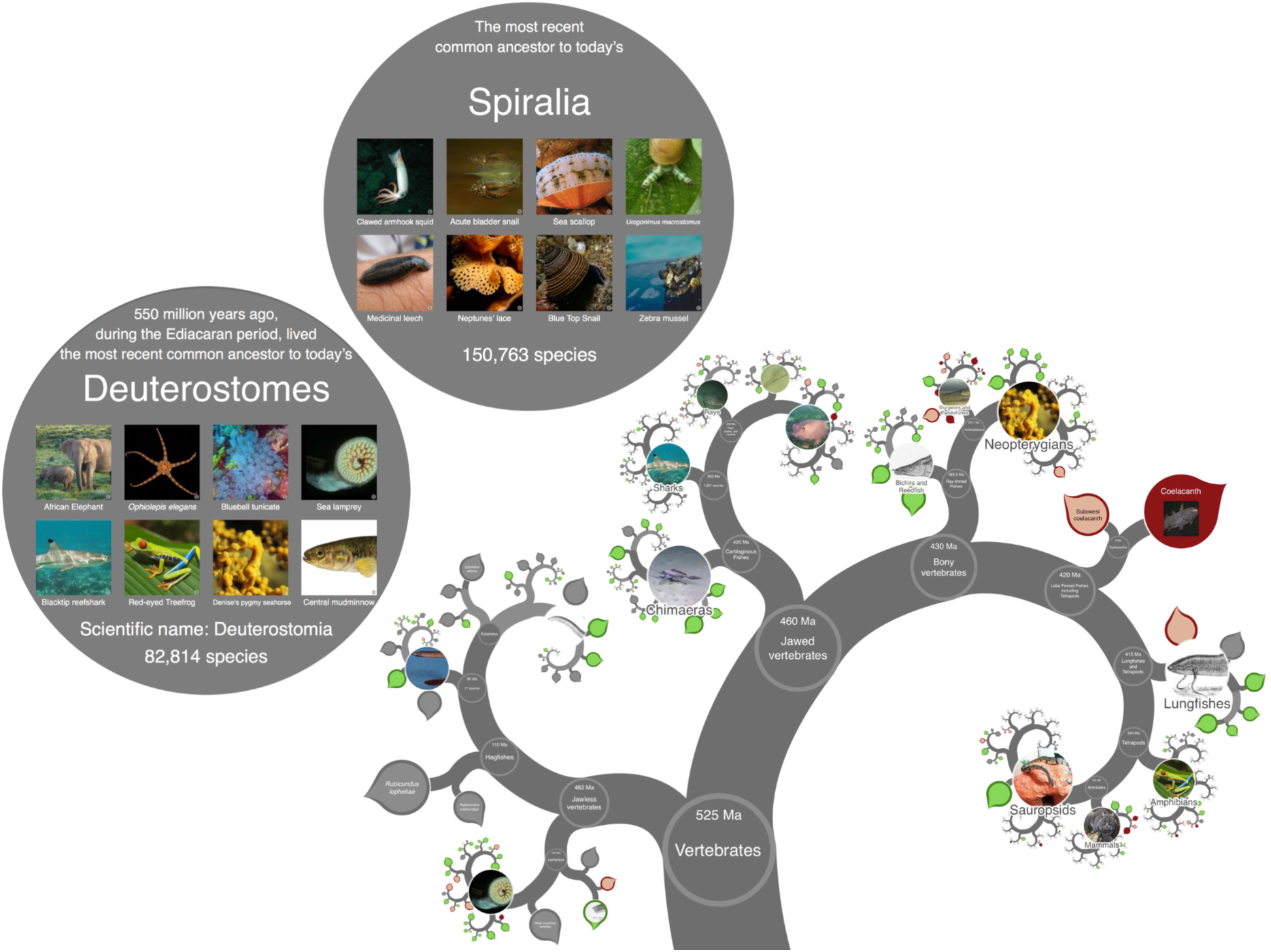
Screenshot of the tree focused on vertebrates composed with expanded views of two interior nodes. The interior nodes (Deuterostomes and Spiralia) each show their eight representative images. All the photographs used are available under a public domain license. Note that an SVG screenshot tool is built into OneZoom, see supporting information S2. The colour scheme for leaves is based on IUCN Red List status of species (see colour key in Figure 5).

### Database

The data overlaid onto the tree is stored in an SQL database. The database contains *leaf* and *node* tables (see Figure 2), with each row corresponding to a taxon with its `ott` ID and associated metadata such as scientific name, ID in other databases and popularity score. Since our ToL explorer uses the compressed tree string to access tree structure, we do not strictly require the database to contain topology information. However, because of the correspondence between taxon position in the string and row number in the tables, the table rows are automatically ordered as a “nested set” (Celko 2004). Accordingly, providing a `leaf_left` and `leaf_right` field for rows in the node table gives rapid access to the set of leaves, and hence the species richness, under any node (they are the leaves whose row numbers lie between these two values). On the same principle, a `node_right` field exists to allow rapid listing of all descendant rows of the node table. For completeness, rows in both tables also have a `parent` field indexing into the node table, and a `real_parent` field giving the closest ancestral row which does not correspond to a “fake” polytomy-resolving node. Additional, optional fields include those for the `age` of a node (or equivalently, `extinction_date`, for a leaf), and in the node table, three sets of fields each containing the OTT ID for the 8 species whose images have been identified by the representative image algorithm.

Additionally, there are tables for vernacular names, thumbnail images, and extinction risk, all of which can be searched for rows corresponding to a given OTT ID, allowing multiple images or multiple vernacular names in different languages to be linked to a single taxon.

### Popularity index algorithm

All species on the OneZoom tree have a “popularity” index, which is used internally in our sponsorship-based funding model (see discussion) but which we make freely available for general use. The popularity of a species is based on sizes and recent views of pages on Wikipedia. We consider Wikipedia a more reliable source of popularity than, for example, search engine hits, as online searches tend to involve vernacular taxon names which not only can have alternative meanings, but are also difficult to map to a specific taxon (see also Correia 2019). Our mapping from OTT ID to Wikidata item (or ‘Q’) ID, can be used to identify titles for taxon pages on Wikipedia; Wikimedia dump files can then be searched for statistics associated with these page titles.

Currently, 307,503 OneZoom taxa have an English-language Wikipedia page. Each is assigned a raw popularity score, constructed by multiplying the average monthly number of visits over the past 2 years (truncated to remove spikes) by the current page size, and taking the square root. Thus pages with a modicum of content and a reasonable number of visits are considered more popular than small pages with many visits or large pages with few visits. Problematically, Wikipedia visitors may not always distinguish species from higher taxonomic levels (e.g. “the honey bee” consists of 8 species). Assigning popularity to individual species, especially those without Wikipedia pages, thus requires us to percolate popularity from higher taxa down to the species level. Similarly, some popular Wikipedia pages (e.g. “dog”) concern subspecific taxa, whose popularity needs to be percolated upward to the containing species. The upward and downward percolation is done by using the phylogenetic tree structure (see supporting information S4): the resulting “phylogenetically-informed popularity metric” can be compared across the tree of life.

### Client-side tree viewer

The basic client-side visualisation code is described in Rosindell and Harmon (2012), here newly modularised and augmented to enable dynamic overlaying of metadata onto the tree. In addition, user controls (zooming, location, search, etc) have become distinct HTML elements which issue commands to an underlying client side *controller* in charge of an HTML5-canvas-based tree drawing engine (see Figure 1). The same controller and drawing engine can thus be embedded into radically different front ends. The original OneZoom project was founded, not based on any one particular tree view, but on the broader idea of fractal algorithms for laying out trees that can be explored with a zooming user interface. A ‘fractal algorithm’ in this context means an approach that compresses the geometry of a tree slightly as we move along branches, enabling more distant sections of the tree to be taken away from the user’s focus; this generates a visualisation with a ‘self similar’ repeating structure indicative of a fractal (Rosindell and Harmon 2012). Within the drawing engine, rendering code is separate from geometric layout calculations – these calculations create what we refer to as a cartographic “projection” from the tree into cartesian space. Because the topology of the tree is stored, users can switch instantly between different projections, an important way to negotiate inevitable trade-offs when visualizing large trees (James Rosindell and Wong 2018). We provide five projections, including a view able to display polytomies with a layout similar to de Vienne (2016). A hierarchical structure of nested Javascript objects represents the tree within the client; during tree exploration this structure is expanded and contracted using the locally stored compressed tree string and list of cutpoints (Figure 2), with all locally expanded nodes of the tree remaining connected to a single root node. The controller maintains a background loop which repeatedly calls the server API to request further metadata for leaves and nodes where required, including scientific and vernacular name, node date, representative images, and IUCN Red List status; these are then injected into the local structure which is repeatedly re-rendered. If a zooming “flight” through the tree is required, a path to the destination as well as a suitable amount of metadata is pre-populated before the flight commences.

Taxonomic ID mapping allows us to reliably identify online resources in other databases. In particular, we use the wikidata QID to access wikipedia pages in multiple languages for over 1.3 million taxa on the tree. This allows the standard front end to display appropriate wikipedia information in a tabbed window within our ToL explorer when leaves or nodes are clicked by the user; other tabs are provided to display pages from alternative online reference sources such as EOL, IUCN, GBIF and NCBI.

## Results

The result of this work is best appreciated by visiting the official site www.onezoom.org (a selection of screenshots are provided in Figure 3). The site has been used by nearly 1.5 million unique users, including members of the public, educators, and students of all ages. OneZoom software has also aided many other scientific, educational, and public outreach projects, as well as being endorsed by a wide variety of different user groups, including prominent members of the scientific community. Interaction with the explorer is smooth even on relatively old mobile devices (e.g. a first generation iPhone SE).

As an example of an alternative front end, we make a freely accessible implementation of OneZoom customised for use in museums and other public locations (see supporting information S2). This has a simplified user interface with labelled buttons, a screensaver to attract walk-by users, an introductory automated tour, and sandboxing functionality so that users cannot follow links to other internet sites, even within Wikipedia popups. This is currently used by the touring “Darwin and Dinosaurs” exhibition, earlier versions have been used around the globe including at the Australian Museum in Sydney. Generalization of the tree layout code has also enabled a range of switchable tree views and colour schemes (see Figure 4 and Figure 5). The view flexibility has in turn enabled third party projects to restyle the tree to fit their own needs. Notably, this includes the One Tree One Planet project (https://www.onetreeoneplanet.org) a digital artwork by the artist Naziha Mestaoui in collaboration with scientists at the Florida Museum of Natural History, aimed at encouraging individuals to take personal actions for biodiversity and conservation.

**Figure 4:**
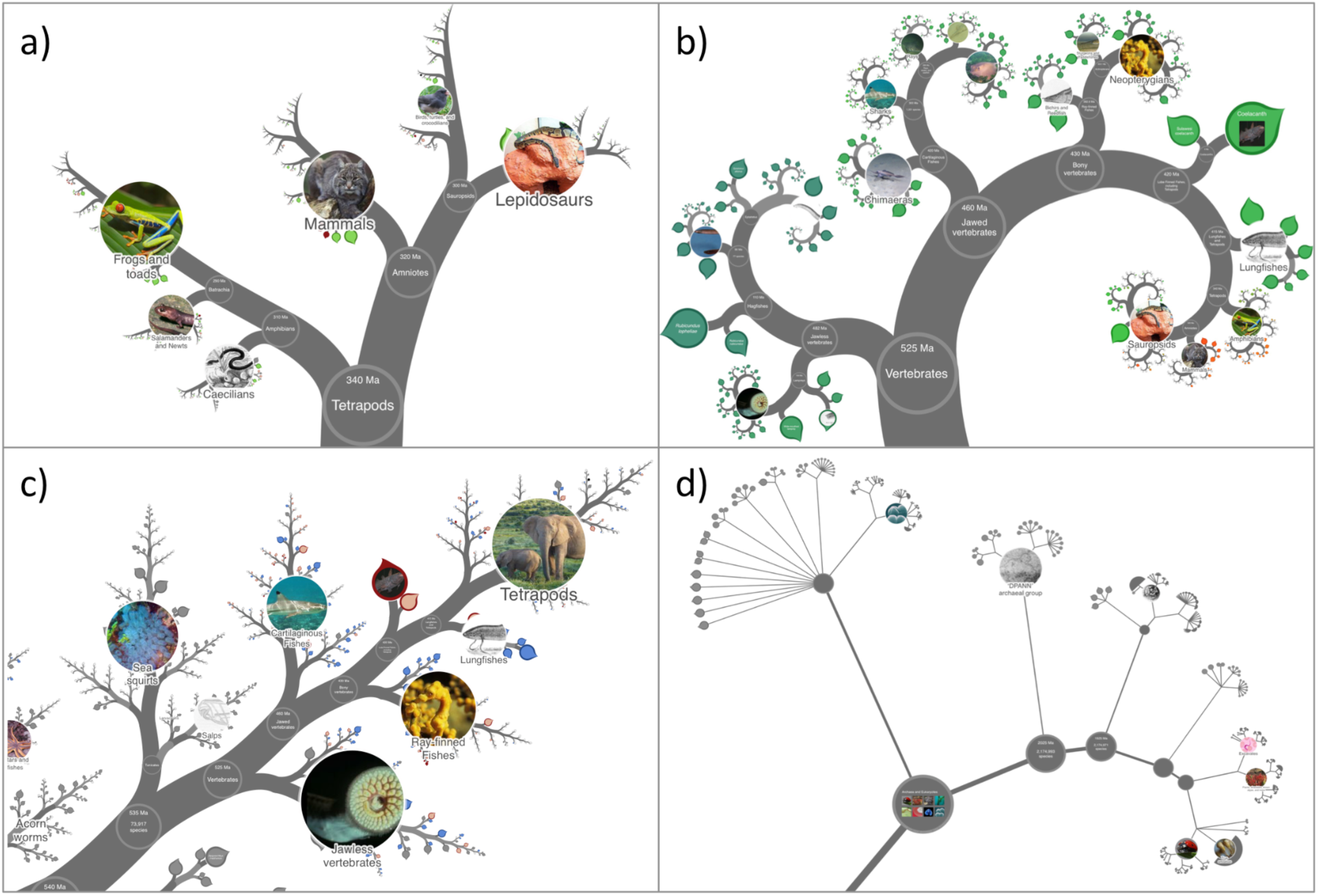
Four different projections implemented on OneZoom. Panel a) “Natural” projection with IUCN Red List colour scheme, panel b) “Spiral” projection with popularity index colour scheme as a gradient from dark blue (least popular) through greens to bright red (most popular), panel c) “Fern” projection with colour blind friendly IUCN Red List colour scheme, panel d) “Polytomy” projection with IUCN Red List colour scheme (see also colour scheme keys in Figure 5).

**Figure 5:**
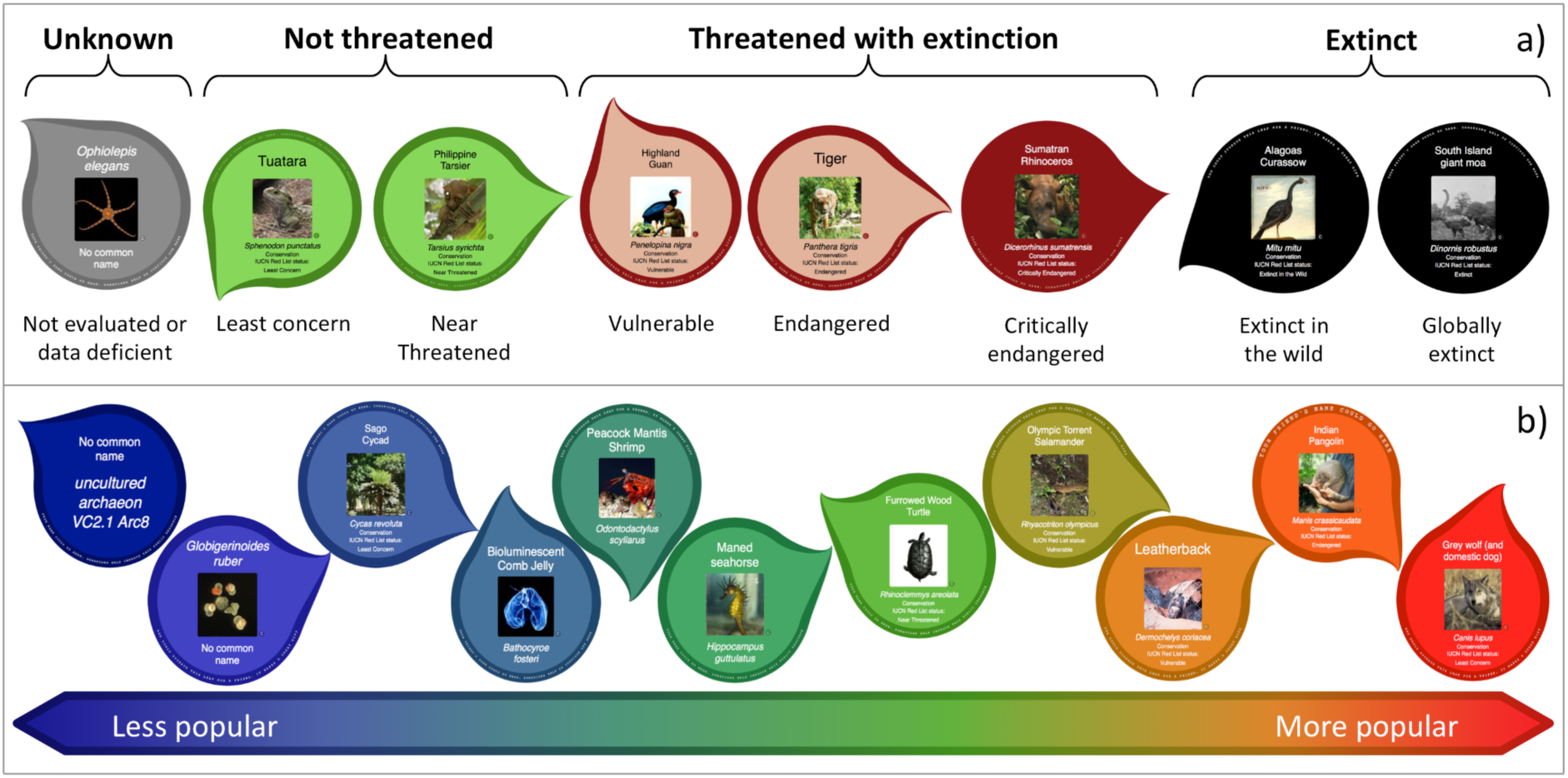
Keys for two main colour schemes based on IUCN Red List status of species (panel a), and based on popularity index as a continuous colour gradient (panel b). We purposefully use a circle for globally extinct species to visually mirror interior nodes of the tree that also represent extinct species of particular relevance.

The popularity index for all species is a work in progress but results are generally consistent with our expectations of popular species (see Figure 6 and Figure 7). It has been made available by public API (see supporting information S1) and has been re-used by the Phylotastic project (Stoltzfus et al. 2013), contributed ID mappings to related work (Millard et al. 2020) and formed the basis for our own novel sustainability model (see discussion).

**Figure 6:**
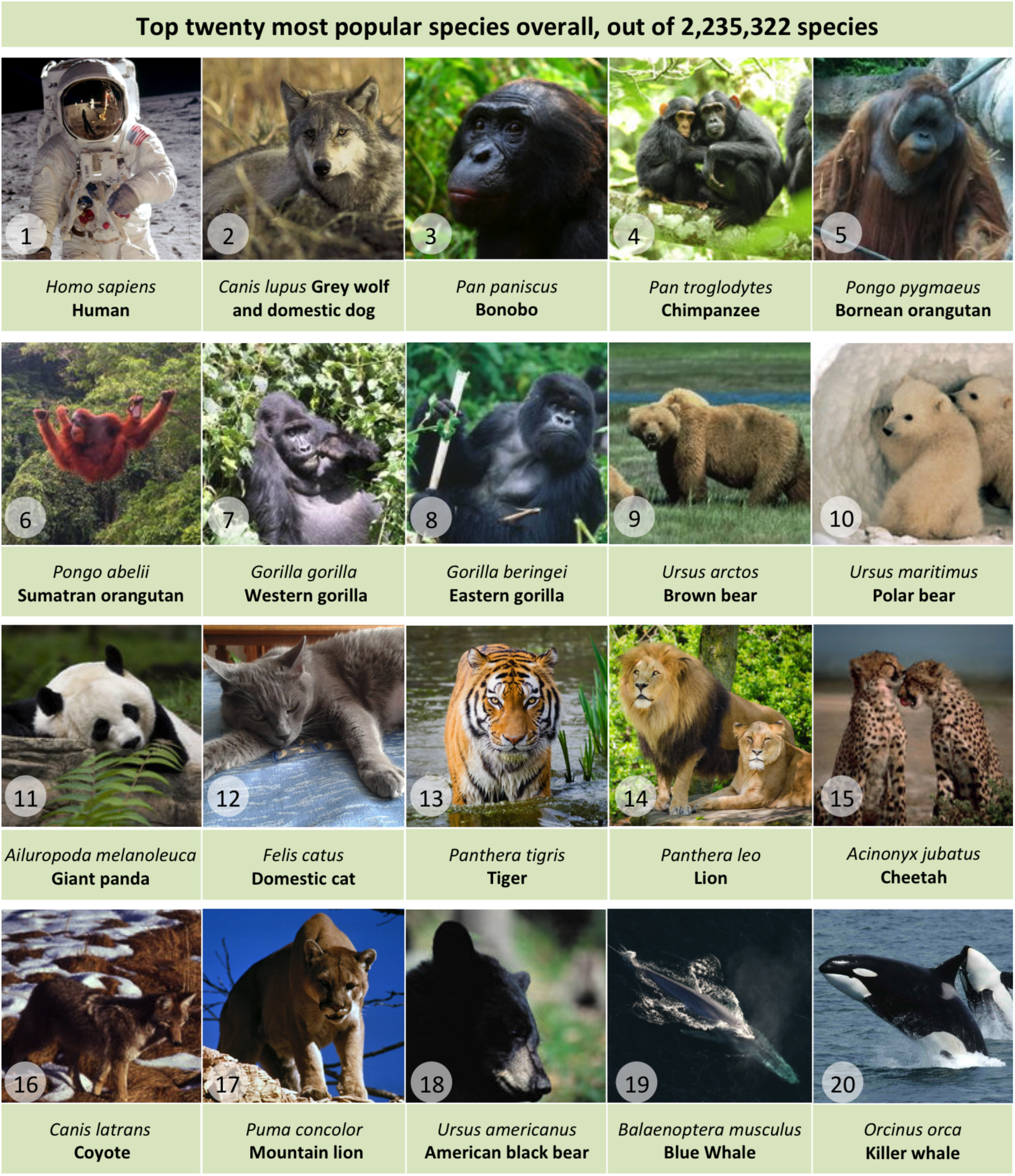
The top twenty most popular species across the entire tree of life, illustrated by public domain images. The most popular species are all charismatic mammals.

**Figure 7:**
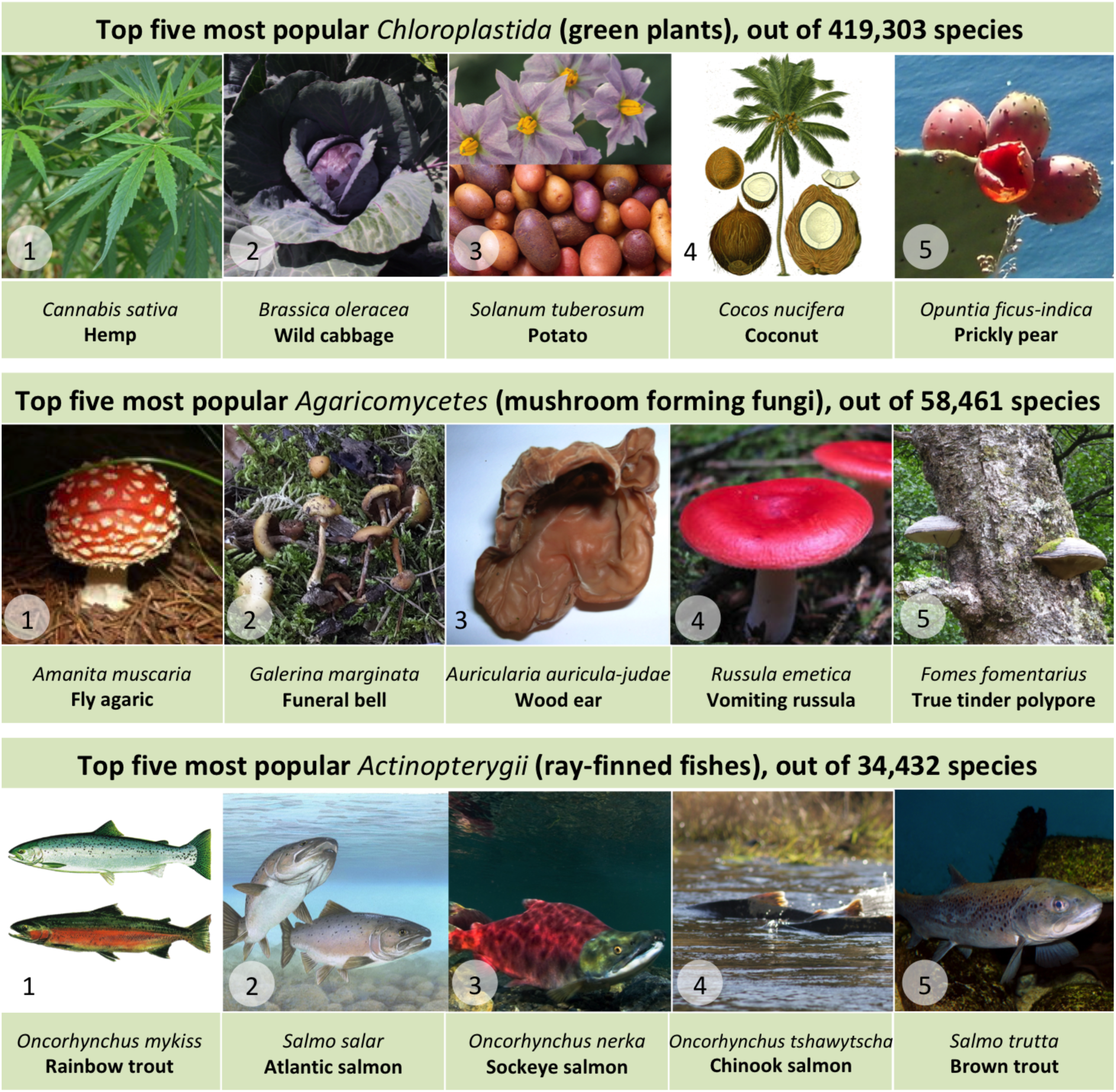
Showing the top five most popular species of green plants, mushroom-forming fungi and ray finned fish. These species are all noteworthy either via use in food or recreation, or for their toxicity.

## Discussion

With this work, we believe that we have realised the original 2012 vision for OneZoom, to create a “Google Earth of Biology”: a complete and engaging tree of life explorer available to all as a community resource. Derived data, accessible via APIs, enable a wide range of work transcending the use-case of a tree of life explorer. For example, images are essential for communicating biodiversity, so our representative image algorithm should find multiple uses, such as decorating taxon information pages, embellishing slides, or in educational games. The popularity index has the potential to inform conservation efforts, especially where evidence of heightened or lessened public interest is required.

We can now reflect on the challenges OneZoom has faced, and suggest next steps. In an attempt to solve the ubiquitous problem of funding and sustaining science projects, OneZoom has registered as an independent Charitable Incorporated Organisation (non-profit), and developed a new, and to the best of our knowledge unique, funding model. Rather than seek income through uncontrolled advertising, we offer visitors the opportunity to “sponsor” leaves on the tree of life for four year blocks of time, with only one name appearing on each leaf. Expected donation amounts for sponsoring increase with the popularity of the species according to our popularity index. Sponsors commonly choose species of personal significance, or sponsor as a gift to others (see examples at http://www.onezoom.org/sponsored). Sponsorship has provided a consistent stream of income sufficient to maintain the website and implement small software improvements, but is not yet sufficient for major development work. We think that addition of sponsorship has also added value to the tree by showing many more ways in which it connects to real people though those who chose to sponsor.

Future work in this area should seek new fractally inspired layout algorithms to represent dated trees and trees with extinct species. Inclusion of extinct species presents specific challenges, both in phylogenetic placement, taxonomy, and visualization; we therefore foresee a need to maintain three trees of scientifically described species: i) all living species, perhaps including only recent extinctions with anthropogenic causes ii) all species, both living and extinct, iii) all living species plus a subset of popular extinct species. OneZoom members have endeavoured to contribute advice and code to other projects, particularly OpenTree and the Encyclopedia of Life, and it is important to maintain this spirit of collaboration: for example we hope in future to improve how data errors reported to us are disseminated back to these data sources. Finally, to engage non-experts, we wish to create a framework for creating engaging “tours” around the tree of life, generated by users and volunteers. These would involve zooming between a series of locations around the tree, each location featuring additional audio, video, or text-based content. We envisage this as a new approach to engage the public in biodiversity, conservation and evolution, within the context of a complete phylogeny of life.

## Author contributions

Both authors wrote and revised the manuscript, J.R. made the figures with feedback from Y.W. Y.W. wrote the popularity index, ID matching algorithms and image downloading scripts. J.R. wrote the representative image selection scripts. Y.W. synthesised the tree. J.R. wrote the tree viewer, which was later refactored by professional developers. All software development was conducted with regular discussion between both authors.

## Acknowledgements

Much of the data for OneZoom comes from the Open Tree of Life and Encyclopedia of Life projects, who we thank for their support, both with data and responses to our specific requests. We also thank Richard Dawkins for his generous financial support and promotion of OneZoom, Jonathan Drori for invaluable advice and generous financial support, Luke Harmon for playing a key role in the early stages of the project and for his contributions as board member of the OneZoom organisation, Jonathan Sutton for contributing to the OneZoom codebase during his summer placement, Jamie Lentin and Kai Zhong for software development work as hired developers, and the many other donors who kindly sponsored leaves on the OneZoom tree of life making it possible to sustain and grow the OneZoom project. We thank Naziha Mestaoui, who sadly passed away in April 2020, for her artistic leadership of the One Tree One Planet project. We thank Douglas Soltis, Pamela Soltis, Robert Guralnick, Matt Gitzendanner for their scientific contributions to the One Tree One Planet project. We thank numerous artists and photographers for their public domain work included in this manuscript. OneZoom CIO received a small payment from University of Maryland, as part of the Phylotastic project, to provide an API to access the popularity index. J.R. was part of an Imperial College London project that received funding from INEOS to work on Atlantic Salmon populations (the second most popular ray finned fish according to the present submission). J.R. was supported by fellowships from NERC (NE/I021179, NE/L011611/1). Through J.R., this study is an output of the Georgina Mace centre for the Living Planet at Imperial College London. J.R. thanks Georgina Mace, who sadly passed away in September 2020, for her early encouragement and support of OneZoom since 2012 when the project was in its infancy.

