## Supplementary information for "Dynamic visualisation of million-tip trees: the OneZoom project"

#### Supporting Information S1: Available OneZoom APIs

Much of the novel data synthesis described in the main text can be accessed via online APIs. In particular, the popularity index is accessible and fully documented at http://www.onezoom.org/popularity. Other public APIs provide access to mappings from OTT IDs to identifiers in other databases, representative thumbnail images for higher taxa, and vernacular names associated with OTT IDs, these are documented at http://www.onezoom.org/API.

#### Supporting Information S2: List of relevant URLs

Here we list key links to online software resources and tools arising from this work.

##### Software

Full OneZoom repository on GitHub: https://github.com/OneZoom/OZtree

Docker image https://hub.docker.com/r/onezoom/oztree-complete

Docker image (reduced file size) https://hub.docker.com/r/onezoom/oztree

Information for developers https://www.onezoom.org/developer.html

Tree building code:
https://github.com/OneZoom/OZtree/tree/master/OZprivate/ServerScripts/TreeBuild

Taxon mapping and popularity code:
https://github.com/OneZoom/OZtree/tree/master/OZprivate/ServerScripts/TaxonMappingAndPopularity

Image processing code:
https://github.com/OneZoom/OZtree/tree/master/OZprivate/ServerScripts/Utilities

Javascript for tree viewer module and associated projections (view types):
https://github.com/OneZoom/OZtree/tree/master/OZprivate/rawJS/OZTreeModule

#### Resources

OneZoom website main landing page: www.onezoom.org

Complete how to use guide https://www.onezoom.org/full_guide.html

Popular species API details: http://www.onezoom.org/popularity/index.html

OneZoom website tree viewer: http://www.onezoom.org/life

OneZoom museum display exhibit software:
http://www.onezoom.org/education/museum_display_setup.html

OneZoom charity (not for profit) organisation details:
http://www.onezoom.org/about.html
http://www.onezoom.org/static/images/RegistrationCertificate.jpg

OneZoom SVG screenshot tool
http://www.onezoom.org/education/screenshot_launcher.html

### Supporting Information S3: Tree construction

We provide a set of open source Python and Perl scripts to augment a nested series of bespoke phylogenetic trees with additional subtrees from the Open Tree of Life (Hinchliff *et al.*, 2015), producing a single 2.2 million tip Newick-format phylogenetic tree. Currently, our bespoke trees are used to provide a robust backbone for the complete OneZoom tree of life. For example, and in contrast to the current OpenTree, this allows us to impose a paraphyletic Archaea (Williams and Embley, 2014), to display particular hypotheses for relationships between the major groups of eukarotes (Burki, 2014), to break polytomies in an informed manner, such as at the root of the placental mammals (Poulakakis and Stamatakis, 2010), and to remove misplaced OpenTree taxa.

Bespoke phylogenies are provided in Newick format, and OTT IDs then associated with named taxa by querying the OpenTree TNRS service against the taxon's scientific name, producing a set of phylogenies with tips labelled, for example, *Homo_sapiens_ott770315*. These trees are then merged together into a single bespoke Newick tree using a simple string substitution process. Bespoke trees may also contain tips that are not species, but instead consist of a name and OTT ID followed by an '@' symbol: these correspond to higher-level taxa, and the OTT IDs are collected and used to extract the matching subtrees from the downloadable Newick representation of the synthetic OpenTree (currently version 13.4). A full tree is then constructed by replacing each such tip in the bespoke Newick tree with its corresponding subtree. Note that unlike the displayed OneZoom phylogeny, the resulting tree may have polytomies, unary nodes, and taxa below the species level (e.g. subspecies). The compressed tree string, which removes these features of the full tree, is created from the full Newick as part of the identifier mapping process (section S4).

The current "AllLife" tree, as displayed on the OneZoom website, is constructed by combining a set of 47 bespoke Newick-format trees, many from a more limited dated synthetic phylogeny described in Dawkins and Wong (2016), with 438 subtrees from the synthetic OpenTree. Dates for nodes on the OneZoom tree are based on a variety of publications, with branch lengths, where provided, adjusted by the PATHd8 software (Britton *et al.*, 2007) to ensure ultrametricity. Details, full academic references, and links to the bespoke trees are provided at http://www.onezoom.org/data_sources.html, and code to generate the tree, along with reproducible instructions, is on the OneZoom GitHub repository, within the OZprivate/ServerScripts/TreeBuild directory. The URL http://www.onezoom.org/static/FinalOutputs/AllLife_full_tree.phy.gz provides a direct download of the current version of the full Newick tree.

### Supporting Information S4: Identifier mapping and popularity calculation

For datasets of millions of taxonomic items, using online APIs is often not feasible. Instead we map identifiers in external datasets by supplementing the OpenTree taxonomy file with database dumps provided by EOL and Wikidata. For popularity scoring, the wikidata identifiers then allow us to match against page size and page views download dumps provided by Wikipedia.

A single Python script is used to (1) map taxonomic identifiers between datasets (2) calculate popularities by matching against wikipedia page data and (3) output files suitable for import into the OneZoom database. The first two steps can be time consuming, largely because of the size of the datasets used (e.g. the uncompressed Wikidata JSON dump is greater than a terabyte in size): total runtime on a 64 CPU Intel Xeon E5 server is roughly 1 hour. Below, we provide an algorithmic overview: more details, together with the open source Python code, are available online at
https://github.com/OneZoom/OZtree/tree/master/OZprivate/ServerScripts/TaxonMappingAndPopularity.

#### Identifier Mapping

Initially, the full OTT-labelled OneZoom Newick tree, produced as per supporting information S3, is read in using version 4 of the Dendropy library (Sukumaran and Holder, 2010). Each node on the tree is treated as a taxon item, and where possible indexed by its OTT ID. The current version of the OpenTree taxonomy file (from https://tree.opentreeoflife.org/about/taxonomy-version/) is then parsed, and the OTT ID index used to associate taxon items with original source identifiers. For example, in OTT version 3.2, the line in the file corresponding to own species (OTT ID: 770315) includes the following:

| uid | parent uid | name | rank | sourceinfo |
|---|---|---|---|---|
| 770315 | 770309 | Homo sapiens | species | ncbi:9606,gbif:2436436,irmng:10857762,irmng:11388931 |

The `sourceinfo` field indicates that the identifier for the species labelled *Homo sapiens* in the OpenTree corresponds to

1. Taxon 9606 in the National Center for Biotechnology Information taxonomic system (e.g. GenBank: see https://www.ncbi.nlm.nih.gov/Taxonomy/Browser/wwwtax.cgi?id=9606)
2. taxon 2436436 in the Global Biodiversity Information Facility taxonomy (GBIF: see https://www.gbif.org/species/2436436)
3. taxa 10857762 and 11388931 in the Internet Register of Marine and Nonmarine Genera (IRMNG: see https://www.irmng.org/aphia.php?p=taxdetails&id=10857762; the second IRMNG taxon corresponds to *Homo floresiensis*, currently treated by the OpenTree as a synonym).

Other common sources in the OpenTree are the World Register of Marine Species (WoRMS) ID, the Index Fungorum ID (for fungi), and the SILVA ribosomal RNA database ID (for bacteria).

Through their OTT ID, taxa in the tree are therefore associated with (1) NCBI, (2) Index Fungorum, (3) WoRMS, (4) IRMNG, and (5) GBIF identifiers. These can then be used to map to equivalent identifiers in the Encyclopedia of Life (the EOL "page_id"), Wikidata (the "Q" ID), and the IUCN (the RedList "species ID"). OneZoom currently uses EOL for images and common names: at our request, EOL now provides a regular identifier map that includes, where available, the NBCI, WoRMS, and GBIF IDs for each EoL page (see https://opendata.eol.org/dataset/identifier-map). Our script matches these EOL-provided IDs against the stored versions in each taxon item. This enables us to match OpenTree taxa to EOL page_ids without the need for explicit taxonomic name resolution. If two or more EOL pages match against a single OpenTree taxon, the match supported by the majority of identifiers is taken; the identifier order listed at the start of this paragraph is used to resolve ties.

A similar procedure is undertaken using the Wikidata JSON download dump (https://dumps.wikimedia.org/wikidatawiki/entities/), focussing on Wikidata items that are an instance of the "taxon" entity (https://www.wikidata.org/wiki/Q16521). These Wikidata "taxon" items commonly include properties corresponding to identifiers from other data sources, including the five described above (Wikidata properties 685, 1391, 850, 5055 and 846, see e.g. https://www.wikidata.org/wiki/Property:P685). These items also contain "sitelinks" to the

Unicode page titles used in various language Wikipedias. We store page titles for one specific language (by default, for the English language Wikipedia), and use a bitfield to record the presence or absence of sitelinks in the 20 most popular language Wikipedias. Where a wikipedia page contains an EOL (P830) or IUCN (P627) property, these are also recorded. Finally, certain popular taxa (e.g. dogs, cows, and humans) have both a taxon item and a Wikidata item corresponding to the common name concept of the taxon (e.g. the taxon item for the domestic dog, *Canis lupus familiaris*, is Q26972265, whereas the concept of a dog is encapsulated by Q144). Although database identifiers are associated with the taxon item, sitelinks to commonly visited wikipedia pages are often contained in the common name concept, which is linked to the original taxon item via the entity "organisms known by a particular common name" (Q55983715: currently there are 48 such items, which are listed at https://w.wiki/enx). During the scan of the Wikidata dump, we also collect these items, to allow for later overwriting of the QID and the sitelinks of the original taxon item with their common name equivalents. The Wikidata dump file is stored in .bzip2 format, which allows independent compressed chunks to be decompressed, parsed, and taxon information extracted in parallel, representing a considerable speed-up on multiprocessor machines (it can take over 8 hours on a modern single processor machine simply to decompress the ~50GB compressed Wikidata dump file).

#### Popularity Index

Once wikipedia page titles have been collected, they can be queried for relevant statistics. Again, due to the number of queries involved, this is most efficiently done using downloaded database dumps. The current size in bytes of the wikitext for a wikipedia page (which does not count transcluded template), is available as one of the fields in the latest-page-sql dump from http://dumps.wikimedia.org/enwiki. Compressed monthly pageviews since 2011 for all wikipedias are available as individual bzip2 compressed files, each roughly half a gigabyte in size, from http://dumps.wikimedia.org/other/pagecounts-ez/. The popularity algorithm extracts information from both of these sources and appends both the current page size and, by default the monthly number of visits for the past 24 months to each of the roughly 0.3 million nodes on the tree which have a corresponding English language Wikipedia page. Since Wikipedia pageviews can experience unusual surges due to the capricious nature of online trends, we calculate an average monthly pageview after removing the two months with the highest visit number: this ensures the removal of a large surge should it cross a monthly boundary.

As described in the main text, a relative popularity measure, the "raw popularity", is calculated by multiplying the page size by the trimmed average monthly pageview, and taking the square root. Most species will not have an associated wikipedia page, and therefore no raw popularity, but it seems reasonable that a butterfly species without a Wikipedia page should be treated as more popular than (say) an equivalently undocumented nematode species. In essence, some of the "popularity" of butterflies in general should, at least partly, be inherited by its constituent members. However, taxonomic approaches are heterogeneous across the complete tree of life with some parts of the tree being much more finely divided into named taxa than others. For example primate genera contain rather few species whose divergence times are measured in a few millions of years, whereas the plant genus *Astragalus* contains over 3000 species and famously, a single brachiopod genus

*Lingula* extends back many hundreds of millions of years. Furthermore, taxonomic levels themselves, genus, family, order etc. are not intended to be comparable at the scale of the complete tree. The ideal phylogenetically informed popularity score should therefore aim to be as invariant as possible to the taxonomic approach. For this reason, we only consider algorithms that make the final popularity score for a given taxon invariant to the exchange of the raw popularity scores between any of the higher taxa to which it belongs. Two natural approaches that each meet this requirement are summing and averaging the raw popularity scores of descendants and ancestors.

The problem now is that taxa may be arbitrarily placed in either few or many nested taxonomic levels. When summing, this may give an unjustified advantage to species that belong to many finely divided taxonomic levels (subfamilies, infrafamilies, tribes, subtribes, etc.). However, some of these minor taxonomic levels, even if they have a Wikipedia pages, may receive few visits, so when averaging would unjustifiably decrease the final popularity. We note that summing would be the correct approach if wikipedia visits among all the descendants of (say) Insecta were unaffected by the number of taxonomic subdivisions within the group. Conversely, averaging would be correct if users methodically explored all taxonomic levels of their taxon of interest, regardless of how many there are. In practice the reality is neither of these, and so as a first attempt to compromise between the two extremes, instead of dividing the sum of ancestor and descendant raw popularities by unity (the sum), or dividing it by the number of popularities, $n$ (the average), our phylogenetic popularity measure is calculated by dividing by the natural logarithm of $n$.

#### Database table output and final processing

Following assignment of popularity metrics to tree nodes, polytomies in the tree are resolved randomly, taxa at the level of subspecies and lower are removed, unifurcating nodes are deleted (keeping the highest named node unless the unifurcation ends in a monotypic species, in which case the lowest named node is kept), and the tree ladderized. The final tree, in our bespoke compressed format (see main text) is then output, as well as csv files of the leaf and node tables (Fig 2). A further script is used to construct the cutpoints files used for indexing into the compressed tree string. Additional database fields such as those storing representative images (supporting information S5) and current IUCN categories, are added directly to the database using stand-alone scripts, listed at

https://github.com/OneZoom/OZtree/tree/master/OZprivate/ServerScripts/Utilities.

### Supporting Information S5: Images and image quality scoring

Previous versions of EOL allowed users to assign a rating score of 1-5 for images and other media items, as well as to select crop areas. Ratings were obtained through the EOL v2 API; they are also available in the current version (3) of EOL although the rating functionality has not yet been reimplemented (J Rice, 2019, pers. comm.). The script to query the API and harvest images is available at https://github.com/OneZoom/OZtree/blob/master/OZprivate/ServerScripts/Utilities/EoLQueryPicsNames.py.

In this script, for each species, the first image returned by the EOL API query is taken as the candidate image for that leaf. This is usually the image with the highest average rating, unless a specific image has been hand picked by a curator as the "exemplar" for that species. We also repeat the EOL query with the restriction that the image must be public domain, and again with the restriction that the image be verified by a curator as correctly identified.

The image URLs returned by these queries are used to download images and crop them to squares (and later usually to circles within the tree viewer). The optimal square crop points obtained from EOL are expanded around a fixed centre until either the edge of the original image is hit, or the image width has been expanded by 12.5%. This expansion avoids cutting off too much of the corners from an ideal square crop, when further cropping to a circle.

#### Representative image algorithm

The first step in choosing representative images for higher taxa is to multiply the average EOL image rating (which ranges from 1-5) by 10,000 and round to obtain an integer score between 10,000 to 50,000; any images with no votes are given a score of 25,000 by default. Users only tend to rate an image as two or less if it is poor quality: images with a score of less than 25,000 are therefore discarded from consideration as representatives of higher taxa. The scores of remaining images still under consideration are then augmented to take into account the number of votes cast by deducting

$$max\left(0,\frac{9000}{8}\cdot\left(8-\sum_{i=1}^{5} V_i\right)\right)$$

from the integer score, where $V_i$ is the number of original EOL user votes that rate the image with a score of i (for $1 \leq i \leq 5$). This formula penalises linearly for having fewer than 8 original EOL ratings contributing to the average rating. To avoid images with low numbers of ratings clustering at discrete locations on the scale, the final scores are then jittered by adding a random number drawn from a normal distribution with mean 0 and variance 100.

The postorder traversal algorithm described in the main text gives a representative set of images for higher taxa. The algorithm works its way up the tree selecting up to 8 optimal images for each node based on the sets of optimal images from its child nodes, and the ratings for those images. Phylogenetic position is initially used to order the images at each node (see main text); this list is then re-ordered pairwise until no image scores 2,500 more than the next highest ranking image.

### Supporting Information S6: Database structure

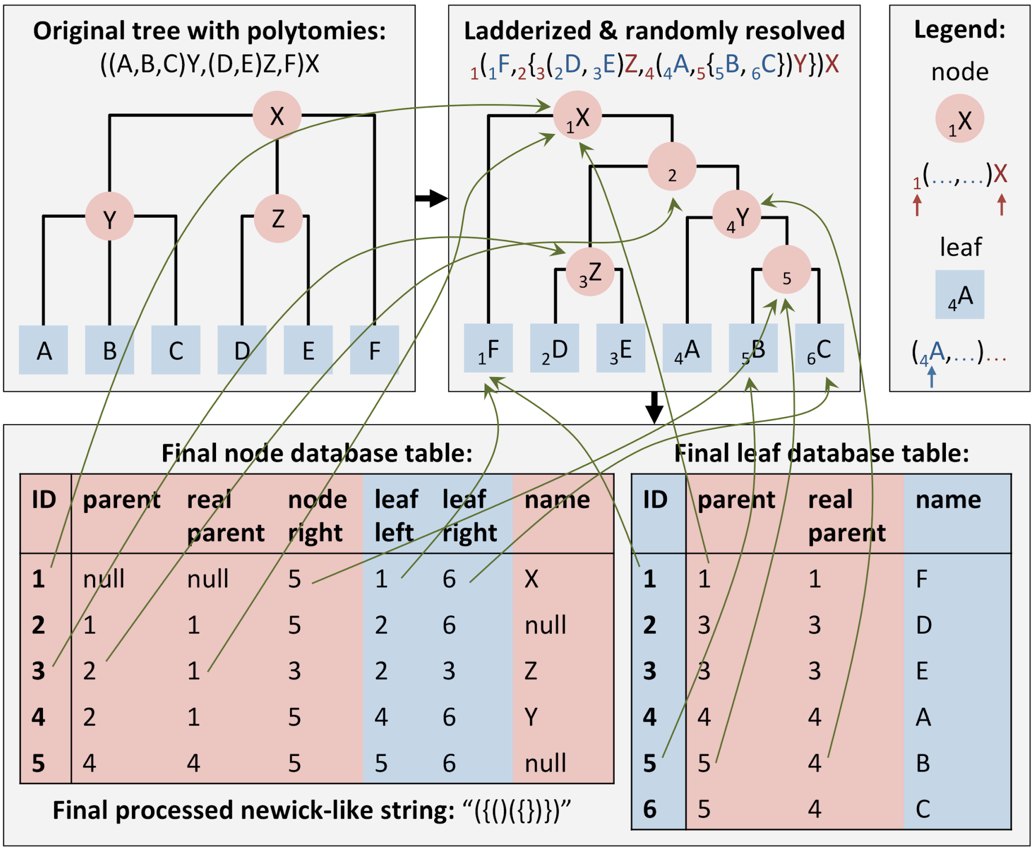


| ID | parent | real parent | node right | leaf left | leaf right | name |
|---|---|---|---|---|---|---|
| 1 | null | null | 5 | 1 | 6 | X |
| 2 | 1 | 1 | 5 | 2 | 6 | null |
| 3 | 2 | 1 | 3 | 2 | 3 | Z |
| 4 | 2 | 1 | 5 | 4 | 6 | Y |
| 5 | 4 | 4 | 5 | 5 | 6 | null |

| ID | parent | real parent | name |
|---|---|---|---|
| 1 | 1 | 1 | F |
| 2 | 3 | 3 | D |
| 3 | 3 | 3 | E |
| 4 | 4 | 4 | A |
| 5 | 5 | 4 | B |
| 6 | 5 | 4 | C |

*Replication of Figure 2 from the main text but with added green arrows to show the source of key numbers in the database tables.*
